# Cortical Entropy Values Correlate with Brain Scale-Free Dynamics

**DOI:** 10.1101/147405

**Authors:** Arturo Tozzi, James F. Peters, Mehmet Niyazi Çankaya

## Abstract

A two-dimensional shadow may encompass more information than its corresponding three-dimensional object. If we rotate the object, we achieve a pool of observed shadows from different angulations, gradients, shapes and variable length contours that make it possible for us to increase our available information. Starting from this simple observation, we show how informational entropies might turn out to be useful in the evaluation of scale-free dynamics in the brain. Indeed, brain activity exhibits a scale-free distribution, which appears as a straight line when plotted in a log power versus log frequency plot. A variation in the scale-free exponent and in the line scaling slope may occur during different functional neurophysiological states. Here we show that modifications in scaling slope are associated with variations in Rényi entropy, a generalization of Shannon informational entropy. From a three-dimensional object’s perspective, by changing its orientation (standing for the cortical scale-free exponent), we detect different two-dimensional shadows from different perception angles (standing for Rènyi entropy in different brain areas). We perform simulations showing how, starting from known values of Rènyi entropy (easily detectable in brain fMRIs or EEG traces), it is feasible to calculate the scaling slope in a given moment and a given brain area. Because changes in scale-free cortical dynamics modify brain activity, suggests the possibility of novel insights in mind reading and description of the forces required for transcranial stimulation.

## INTRODUCTION

Counter-intuitively, a 3D object might encompass less information that a 2D object (we do not see the hidden side of the moon). When observed from a given standpoint, an opaque 3D object could be less informative than a series of its 2D shadows (**Figure 1A**). Here we ask: is there a way to increase the information content of the shadow, to make it possible detect more details about the object, let us say a toy? The answer is positive, if we rotate the toy clockwise (or anticlockwise) along its central axis (**Figures 1B, 1C**). In this case, we achieve a pool of different shadows and shape contours with different angulations, able to increase our total available information. Therefore, 3D object shadows (taken in its totality) are more informative, for a watching observer, whenever he moves. Or, in other words, one could extract more information from a dynamical system, than from a static one. This “toy” analogy gives us the possibility to assess brain function in a novel way. For the purpose of this paper, the toy at rest stands for a given brain scale-free exponent, and its shadows for the corresponding Shannon entropy. In turn, the changes in location of the moving toy stand for changes in brain power law exponents, while the modifications in shadow’s shape and its undulating contour stand for the corresponding Rènyi entropy (Rènyi 1961; Rènyi 1966; Peters 2017; Peters-Ramanna 2016).

**Figure 1.**
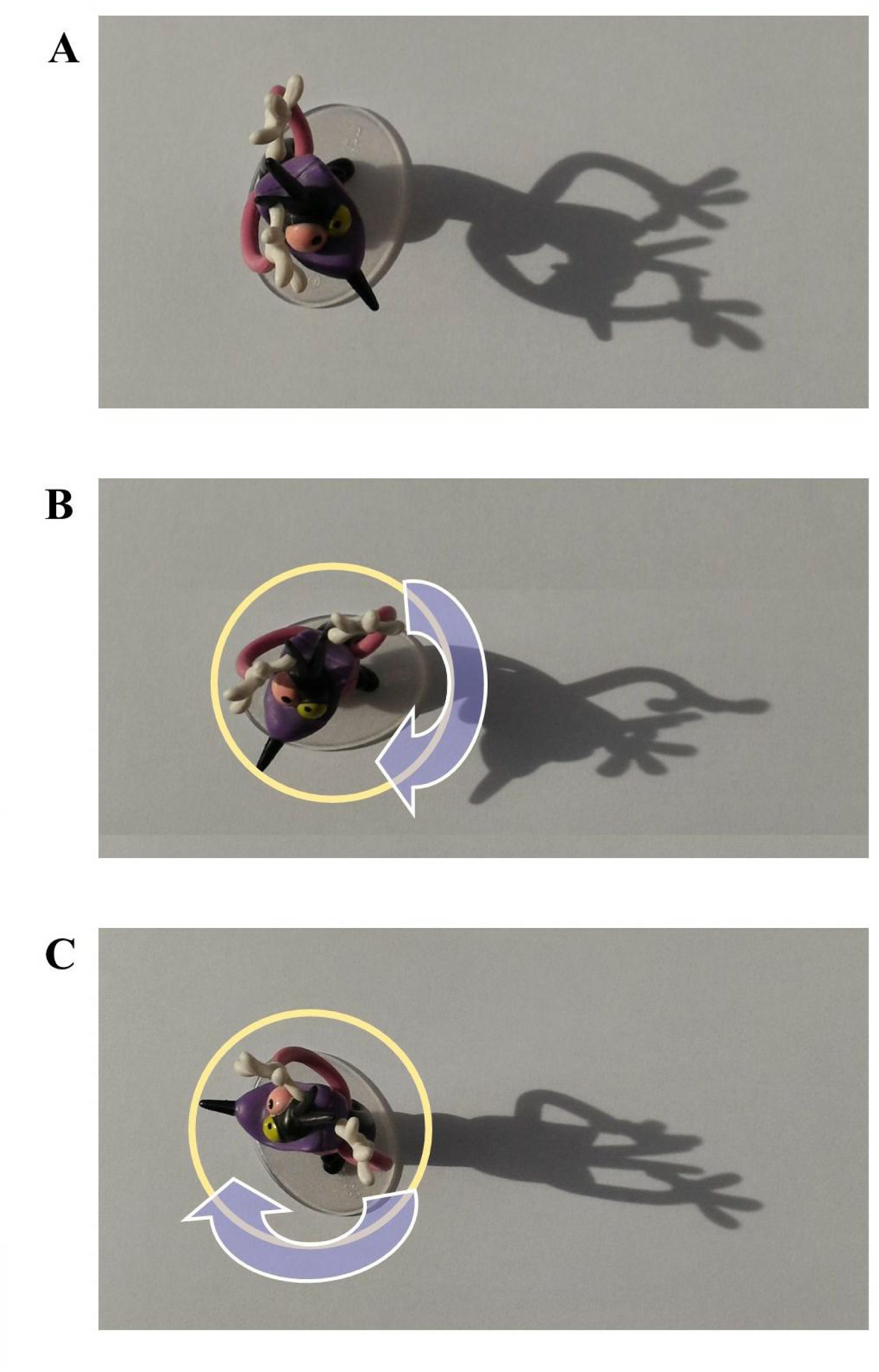
A toy under a light source. In **Figure 1A**, the 2D shadow offers more details of the 3D object than the view on the toy from above. **Figures 1B** and **1C** show how toy rotations lead to shadows with different shapes, and, consequently, with different information content. Therefore, a complete 360 degrees rotation of the object allows a complete evaluation of the shadows’ shapes.

Using our example of a toy as a 3D object and its moving shadows, the aim of this paper is to extract information about such 3D toy, by looking at its moving shadows: in neurophysiological terms, we want to assess the relationships among power-law exponents and generalized informational entropies. Furthermore, we would consider the recent advancement in the neurophenomenology research that established a link between a nested operational architectonics hierarchy of brain functioning (indexed by the EEG field) and the phenomenal (mental) level of brain organization (Fingelkurts and Fingelkurts, 2001; Fingelkurts et al., 2009, 2010, 2013a). According to this line of research, the meanings, which subjectively (mentally) are experienced as thoughts or perceptions, can best be described objectively as created and carried by dynamic fields of neural activity of different sizes that are nested within the operational architectonics hierarchy that organize hundreds of millions of neurons and trillions of synapses (Fingelkurts and Fingelkurts, 2001; Fingelkurts et al., 2009, 2010, 2013a). Starting from Shannon and Rényi entropies, and introducing a link with the scale-free cortical behavior, we show how the brain could be able to change its scaling slopes, in order to control different mental states. We perform simulations in order to demonstrate how, starting from the Rènyi entropy easily detectable in neurodata series (e.g., from brain fMRI or EEG), it is feasible to calculate the cortical scale-free exponent in a given moment, and, consequently, the corresponding mental state.

## MATERIALS AND METHODS

### Shannon entropy on a toy

The brain activity, e.g., in our analogy toy and its movements, can be described in terms of scale-free dynamics. The brain activity observed at many spatiotemporal scales exhibits fluctuations with complex scaling behavior (Newman, 2005; Fingelkurts et al., 2010, 2013a), including not just cortical electric oscillations, but also membrane potentials and neurotransmitter release (Linkenkaer-Hansen et al., 2001; Fox and Raichle, 2007; Milstein et al., 2009). In particular, the frequency spectrum of the cerebral electric activity displays a scale-invariant behaviour, characterized by a given power spectrum, a given frequency and a given exponent, that equals the negative slope of the line in a log power versus log frequency plot (Pritchard, 1992; Van de Ville et al., 2010). In simple terms, a fractal slope 1= stands for a horizontal line parallel to the x axis: the steeper the line, the higher the power law. The (spatial) fractals and (temporal) power laws can be regarded as intrinsic properties of the brain and characterize a large class of neuronal processes (de Arcangelis and Herrmann, 2010; He et al., 2010); moreover, pink noise distributions contain information about how large-scale physiological and pathological outcomes arise from the interactions of many small-scale processes (Jirsa et al., 2014).

Keeping this in mind, a given slope in a given moment of brain activity stands, in our example, for the toy at rest. Toy features can be described through their shadow, e.g., the Shannon entropy. The classical informational Shannon entropy is (Shannon 1948):

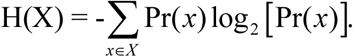

The term X is a random variable with *n* possible outcomes, while *p_i_* = *Pr(x)*, for *i*= 1, 2, 3, …*n*, is a probability distribution *Pr* on a finite set. Shannon entropy states that, under ergodic conditions, if we know the values of *Pr(x)*, we may obtain the values of H(X). In other words, H(X) is a probability density function that defines a generic probability distribution *Pr* such that, if we modify *Pr*, we obtain a different value of entropy on the Shannon’s curve. Hence, changing the probabilities usually occurs when we have prior information. In terms of the object depicted in **Figure 1A**, the steady toy stands for the systems’ microstates, undetectable by macroscopic observers, while the shadow stands for the lower-dimensional Shannon entropy. Looking at the shadow (the Shannon entropy), we achieve otherwise undetectable information about opaque 3D objects such as the toy features.

Now we need to find a brain functional counterpart for the moving toy and its changing shape and varying contours of shadows. Indeed, the scale-free slopes are not constant in the brain dynamics, rather they are characterized by multiple possible exponents during various functional states. It has been demonstrated that different functional states - spontaneous fluctuations, task-evoked, perceptual and motor activity (Buszaki and Watson, 2012), cognitive demands (Buiatti et al., 2007; Fetterhoff et al., 2014), ageing (Suckling et al., 2008) - account for variations in power law exponents across cortical regions (Tinker and Velazquez, 2014; Wink et al., 2008). Accordingly, we may view brain activity as an ensemble of intertwined (mono)fractals, each with its own power slope and scaling range, each marking dynamical transitions between different response regimes (Papo, 2014; see also review Fingelkurts et al., 2013a). In terms of our toy, the whole brain activity stands for all the possible 360 degrees of rotation and their corresponding shadows.

### Introducing Renyi entropy

The 360 degrees changes in toy angulation that gives rise to different shadows can be described in terms of Shannon entropy generalizations. Among the available ones (e.g., Tsallis, 1988), here we favour the one-parameter class of Rényi entropies (Rényi, 1966), a flexible and underestimated information-theory based index (Cambell, 1965). Rényi entropy (1961,1966) is characterized by a changing parameter a andcan be defined as:

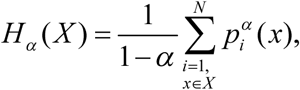

Where 0 ≤ *α* < *∞* and 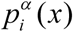 is the probability of event x. The Rényi entropy approaches the Shannon entropy as *α* approaches 1. In other words, the Shannon entropy is a case of Rényi entropy in which α= 1. Rényi entropy has applications in dynamical systems (Hentschel and Proccacia, 1983), coding (Cambell, 1965), information transfer (Jizba et al., 2012), theories in quantum mechanics (Jizba et al., 2015), black holes’ mutual information (Dong, 2016), assessment of human heartbeat fluctuations and coding/noncoding DNA sequences analysis (Costa et al., 2005). Rényi entropy has also been used to quantify ecosystems dynamics and diversity: from land cover types (Carranza et al., 2007; De Luca et al., 2011), to coastal dunes environments (Drius et al., 2013), from urban mosaics (Carranza et al., 2007) to species diversity in large areas (Rocchini et al., 2013).

Unlike Shannon entropy, Rényi entropy makes it possible to describe the system’s status not just at a specific moment, but also when its trend *varies with time* (Müller et al., 2000; Patil and Taillie, 2001). Diversity cannot be reduced to a single index information, since all its aspects cannot be captured in a single statistic (Gorelick, 2006). A complete characterization of landscape diversity can be achieved if, instead of a single index, a parametric family of indices is used, whose members have varying sensitivities to the presence of rare and abundant elements (Jost, 2007). Diversity profiles are flat when the landscape is even, and steeply decrease as the landscape turns uneven (Jost, 2010). Therefore, Rényi’s formulation allows a continuum of possible diversity measures which differ in their sensitivity to the rare and abundant elements in a landscape. Because of its build-in pre-disposition to account for self-similar systems, Rényi entropy is an effective tool to describe multifractal systems (Jizba and Arimitsu, 2001). This means that the generalized fractal dimensional and the Rényi exponent can be thought of as interchangeable. In technical terms, the Rényi parameter is connected via a Legendre transformation with the multifractal singularity spectrum. See Jizba and Arimitsu (2001) and Jizba and Korbel (2014) for technical details on mathematical proof. In particular, this means that Rényi entropy and scale-free exponent (and therefore the scaling slope) are strictly correlated. Therefore, changes in scaling slope lead to changes in the Rényi parameter, and vice versa (Słomczynski et al., 2000): scale-free systems lead to different probability outcomes (in our case, mental functions), based solely on increases or decreases of scale-free exponents. In sum, the “probabilistic” virtues of Rényi entropy represent a novel physics-based approach to probe brain scale-free dynamics, with the potential to lead to new insights into brain systems at all space-time scales and all levels of complexity.

The crucial question, is: What for? How might such a scheme help, in the experimental assessment of brain dynamics? Can the variations (fluctuations) in Rényi entropy can be correlated with the values of a circle perimeter (standing for 360 degrees angles of our toy example rotations)? Our moving toy might help to elucidate such concept. Indeed, toy rotations form an angle on a 360 degrees circumference, so that every angle corresponds to a different shadow, i.e., to a different value of Rényi entropy (Figures 1B and 1C). In neurophysiological terms, we are allowed to calculate a full spectrum of Rènyi entropies, just by evaluating the changes in fractal slopes. The procedure will be described in the next paragraph.

## RESULTS

Here we show the procedure in order to correlate our formulation of Rényi entropy with brain scale-free dynamics. We performed simulations of randomly generated angles on a 2D circle. Every angle displays a random degree, so that their total sum stands for a 360 degrees full circle. The number of rays originating from the circle’s center can be modelled through the cosine theorem, which states that:

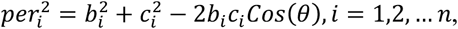

where *b* and *c* (which could be randomly generated in simulations) are the length of rays originating from the circle’s central point. The parameter *per* is one of the three sides of the triangle lying on the circle perimeter (**Figure 2, left side**). The randomly generated parameter *θ* stands for the angle between two rays, while *n* stands for the number of rays. The sum of *per_i_*, namely 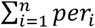, can be regarded as the circle’s perimeter. Each set of perimeter values produces just one Rènyi entropy value: therefore, we replicated the simulation one hundred times.

**Figure 2.**
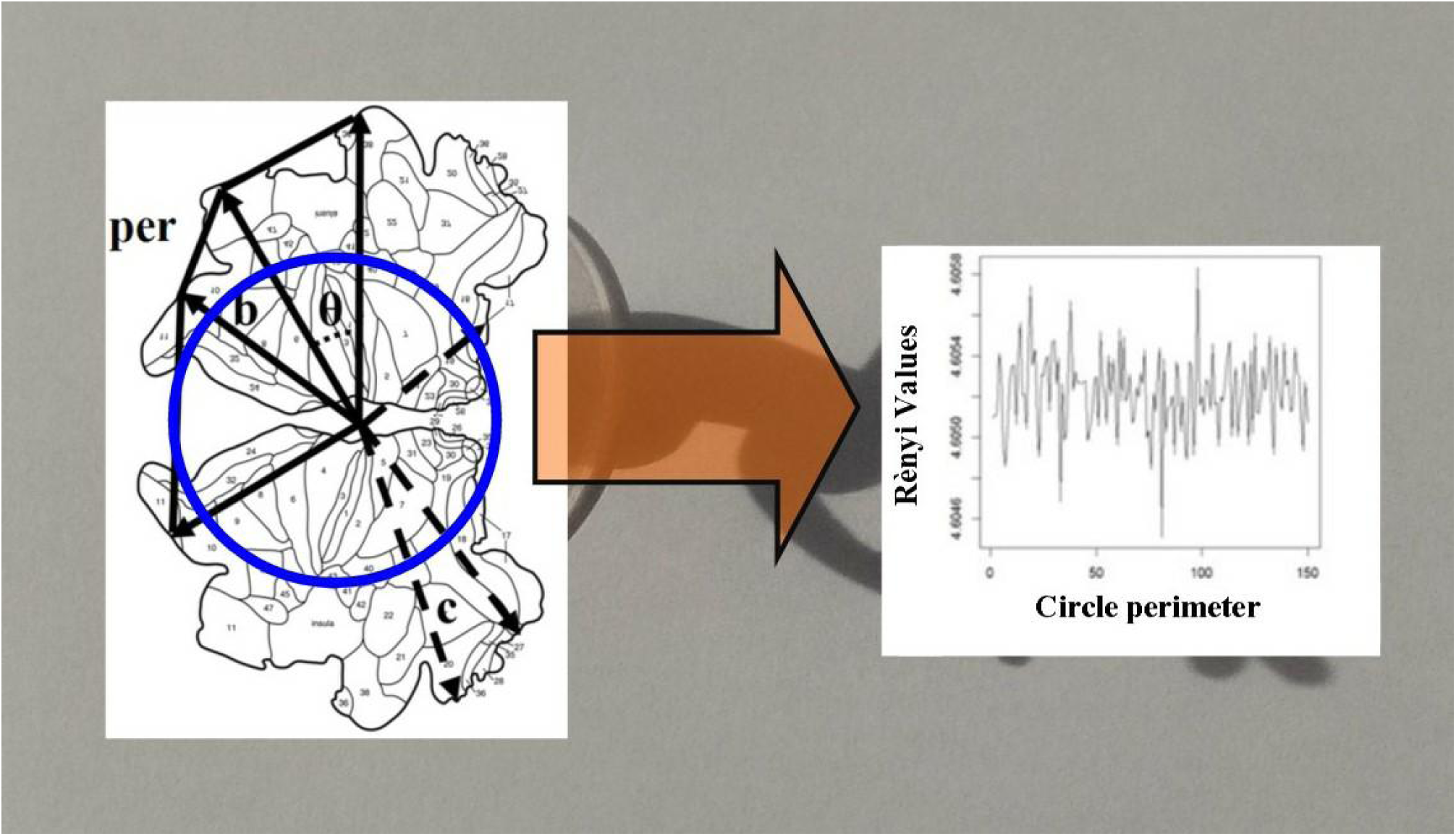
Renyi entropy’ circle perimeter and its values (computed by the Renyi.z function in version 3.2.3 of R software) are superimposed, respectively, to the moving toy and its shadow. **Left part**: Cosine theorem for the evaluation of travel length Rényi entropies in a model of two-dimensional brain (modified from Van Essen, 2005). **Right part:** Simulated Rényi values for each circle’s perimeter values for n=60 and a Rényi exponent=4; θ is generated from a uniform distribution in (0,1) and multiplied by 360. The Rènyi entropy values’ bandwidth is around 4.6.

Note that, in our simulation, the fluctuations in Rényi entropy values depend just on the different perimeter values. In terms of brain assessment, we, instead of evaluating random number, assess the cortical scale-free activity and the scaling slopes. In such a vein, two cerebral hemispheres can be unfolded and flattened into a two-dimensional reconstruction by computerized procedures (Van Essen, 2005). In this model, the nervous electric activity occurs on a 2D brain surface with an approximated circular shape (**Figure 2, left side**). Perimeter values allow us to detect the nervous activity’s travel lengths on the brain surface. After the brain signal’s travel length has been achieved, the corresponding Rènyi Entropy values can be computed (“EntropyEstimation” package: https://cran.r-project.org/web/packages/EntropyEstimation/EntropyEstimation.pdf). **Figure 2, right side,** displays simulated Rényi values obtained from the perimeter values computed through the cosine theorem. In toy’s terms, this means that a larger rotation gives rise to higher levels of information detected from the shadows, while, in brain’s terms, the higher the variations in scaling slopes, the higher the detected Rényi entropy. And vice versa.

## CONCLUSIONS

We described how primitive changes in a single cortical parameter (the scaling slope) may lead to different probability distributions in the brain. We showed that a clear correlation does exist between multifractal spectrum and brain spikes evaluated in terms of probability states. Our novel “inverse”, top-down entropic approach (enabling from known entropy changes, to achieve the unknown probability distributions) allows us to evaluate the macro-states of the nervous system based on a sole order parameter, although we lack a perfect knowledge of micro-states. In our simulations, random numbers generated from different distributions are able to modify the values of Rényi entropy: this means that the brain scale-free spectrum can be extrapolated, just starting from the values of entropy experimentally detected in EEG or fMRI traces. Considering the connection between brain activity dynamics and the mental states (Fingelkurts and Fingelkurts, 2001; Fingelkurts et al., 2009, 2010, 2013a), this observation paves the way to tackle the difficult issue of mind reading.

We hypothesize that, in order to optimize its functions, the brain could be equipped with intrinsic mechanisms of scaling features: in this framework, changes in scale-free exponents play a crucial role in information processing, leading to variations in entropy and probability of mental states. Empirical evidence suggests that cognitive tasks are modulated by the scale-free exponents of the brain fluctuation probability function, leading to a shrinking of multifractal spectrum and/or transitions from mono- to multi-fractal distributions (Popivanov et al., 2006; Fraiman and Chialvo, 2012).

Our results pave the way for innovative diagnostic and therapeutic strategies. For example, the widely-diffused method of pairwise entropy in neuroimaging techniques (Watanabe et al., 2014) might benefit of a reverse Rényi entropy-based approach. It is indeed feasible to start from known functional changes in thermodynamic parameters and/or multifractal slopes, in order to achieve the unknown parameters of probability distribution (the spike frequency). In particular, a change in Rényi exponent, caused by variations in scaling slopes, might lead to different probability outcomes, which can be calculated if just their pairwise entropies are known. Rényi entropy offers a “continuum of possible diversity measures” (Ricotta et al., 2003) at diverse spatiotemporal scales, becoming increasingly regulated by the commonest when α gets higher. The change in α exponent can be regarded as a scaling operation that takes place not in the real, but in the data space (Podani, 1992). We are thus allowed to evaluate how changes of Rényi parameter influence the structure of information measures in the probability space of the scale-free dynamics.

We also conjecture that the scale-free-like brain electric activity - setting aside their supposed relationships with self-organized criticality (Bak et al., 1987), nonequilibrium steady-state dynamics or second order phase transitions (Papo, 2014) - could be modulated through the superimposition of an external electric currents characterized by carefully chosen power law exponents. This may have a practical application. For example, our results suggest that transcranial electrical stimulation’s techniques need to take into account not only the amplitude and frequency of the applied waveforms (Reato et al., 2013), but also their scaling slope. A further tenable field of application are the diseases that have been linked to disturbance of brain networks - Alzheimer’s disease, depression, attention deficit hyperactivity disorder, schizophrenia, autism (Fox and Raichle, 2007; Fingelkurts and Fingelkurts, 2010) -, meaning that they could be ameliorated, or even treated, by appropriate exogenous electromagnetic fields - e.g., via selective application of electric/magnetic waves of specific scale-free exponent on target micro-areas - able to “recover” and restore the physiological brain functions (Sunderam et al., 2009; Fingelkurts and Fingelkurts, 2010, 2015) and related subjective experiences (Fingelkurts et al., 2013b).

